# Colloidal quantum dots suitability for long term cell imaging

**DOI:** 10.1101/425850

**Authors:** Patricia M. A. Farias, André Galembeck, Raquel Milani, Wilson S. Mendonca, Andreas Stingl

## Abstract

Fluorescent semiconductor nanoparticles in tree-dimensional quantum confinement, quantum dots (QDs), synthesized in aqueous medium, and functionalized with polyethylene glycol, were used as probes for the long-term imaging of glial cells. *In vitro* living healthy as well as cancer glial cells were labelled by direct insertion of a small volume of QDs contained in aqueous suspension into the culture wells. A long-term monitoring (over 7 days) of the cells was performed and no evidence of cell fixation and/or damage was observed. Two control groups, healthy and cancer glial cells, were used to compare cell viability. During the observation period, labelled and non labelled cells presented the same dynamics and no difference was observed regarding cell viability. To our knowledge, this is the first report of the viability of hydrophilic prepared quantum dots without any further surface treatment than the polyethylene-glycol coverage for the long-term imaging of living cells. Further, the study also permitted the observation of two distinct interaction mechanisms between cells and QDs. Healthy glial cells were mainly labelled at their surface, while non-healthy glial cells have shown a high rate in the uptake of QDs.

## 1 Introduction

When light interacts with semiconductor nanoparticles, valence band electrons may acquire enough energy to bridge the energy gap and reach the conduction band. When these semiconductor nanoparticles are in the size range of few nanometers (normally 2 up to 10), the electrons small In these cases, the electrons of each particle are confined in each one of the 3 dimensions. Nanoparticles under such conditions constitute examples of low dimensional systems. In such a case, a class of zero dimensional (0D) materials under quantum confinement regime in all the 3 dimensions. These systems are named quantum dots (QDs) [5]. The excited electrons may decay to their original state by means of radiative and/or non-radiative mechanisms [1,2].The radiative ones gave rise to many fluorescence-based assays, while the non-radiative have been used increasingly in photoacoustic (PA) based assays [3,4].

*In vivo* and *in vitro* toxicological studies of QDs have been carried out during the last decade [6-10]. For *in vitro* diagnostic assays, such as fluorescence microscopy (single and multi-photon), QDs have shown to be very efficient, even surpassing conventional dyes, once QDs allow stoichiometrically prepared surfaces for specific targeting in cells and tissue samples, without the need of fixation of such samples. This feature opens entirely new possibilities for bio-imaging based diagnostics techniques. To qualify as probes for in vitro diagnostics, QDs must be biocompatible. This condition may be achieved, both for bottom-up [11-13] and top-down [14-16] prepared QDs, by modifying their surface *via* the controlled addition of layers of organic polyelectrolytes, peptides and/or other biomolecules, such as mono or polyclonal antibodies [17]. Previous work [18] have reported the use of non-polar quantum dots, rendered biocompatible by means of DHLA (dihydrolipoic acid) coverage, to investigate cell uptake/endocytic routes, and conjugated to avidin in order to interact with biotinylated cell surface. In both applications, the QDs were modified in order to remove non-polar functional groups and make them feasible for interacting with biological samples. However, assembly of biomolecule-QD constructs has proven to be a challenge, which still demands several experimental steps [19]. Besides the efficient use of simply produced QDs, the labelling of living cells as performed in this work, potentially may be used as a model for a better understanding of crucial factors that underly the aggressive nature of gliomas [20] and their infiltration in central nervous system.

## 2 Methodology

Otherwise mentioned, chemicals were purchased from Sigma-Aldrich and used without further purification. The C6 rat glioma cells and the healthy glial cells (rat astrocytes) were purchased from Sigma-Aldrich for research purposes only, and are part of the European Collection of Authenticated Cell Culture (ECACC), as declared by Sigma-Aldrich. This way, ethical issues are covered by the certification of ECACC. Core-shell Cadmium Sulphide/Cadmium Hydroxide CdS/Cd(OH)_2_ QDs (4nDOT500) designed for a peak emission of 500 nm were produced by Phornano Holding GmbH, Korneuburg, Austria. The passivation step of CdS nanoparticles consisted of a controlled precipitation of Cd(OH)_2_ layers at high pH medium. QDs were characterized by electronic absorption and emission spectroscopies, conventional and high-resolution transmission electronic microscopy (HR-TEM and TEM) and X-ray diffractometry (XRD). 4nDOT500 are functionalized by polyethylene-glycol (PEG) (molecular weight: 400). QDs were chemically associated to PEG in aqueous media using 0.1M PEG aqueous solution (0.0025 g/mL) at room temperature (RT) at pH 7.2. QDs were centrifuged and two main sizes population were identified: 6 and 9 nm. Rat C6 glioma cells were kept in Dulbecco’s modified Eagle’s medium (DMEM) containing 10% fetal bovine serum (FBS) and 1% penicillin-streptomycin at 37°C in a 5% CO_2_ atmosphere. The C6 cells were seeded at 5×104 cells/well in two sets of 96-well microplates containing DMEM. The cell sets were then named as *L* and *C* (*L* = *Labelled* and *C* = *Control)*. Both sets *L* and *C* sets were analyzed within the same acquisition parameters. Prior to the visualization at the confocal fluorescence microscope set-up, the cells of *L* set were washed with PBS and then, directly visualized by laser scanning confocal microscopy (Leica Microsystems GmbH, Wetzlar, Germany) with an apochromatic water-immersion objective (100X, NA 1.2) at wavelengths of 488 and 533 nm. Fluorescence images presented in next section are the ones resulting from overlaying images obtained by exciting samples at the two wavelengths mentioned above (488 and 533 nm). The same procedure was used for the analysis of control cells (set *C*). Confocal microscopy and viability assays were performed at different intervals during 7 days in order to monitor long-term cell viability and to detect possible morphological damage of the cells. All images were taken with the same acquisition parameters and processed using modular imaging software for fluorescence imaging (Leica QWin Lite, Leica Microsystems GmbH, Wetzlar, Germany) for performing all the assays. Cell viability was then determined using an EZ-Cytox Cell Viability Assay kit according to the manufacturer’s protocol.

The same core/shell Cadmium Sulfide/Cadmium Hydroxide QDs in aqueous medium used in this work, have been previously analyzed by means of *in vivo* toxicological studies using male Wistar rats, which were head-nose exposed to a liquid aerosol of the Cd-based QDs, 6 hours per day on 5 consecutive days [21,22].

## 3 Results

Emission spectrum obtained for as prepared QDs/PEG is presented in Figure 1 (bottom). The maximum emission intensity occurs at 504 nm. In Figure 1 (top), the experimental and the theoretical (vertical lines) X-ray Diffractogram are shown. By analyzing XRD it can be observed that the CdS/Cd(OH)_2_ QDs present cubic crystalline structure, which corresponds to the more stable crystalline form for CdS [23].

**Figure 1-Top:**
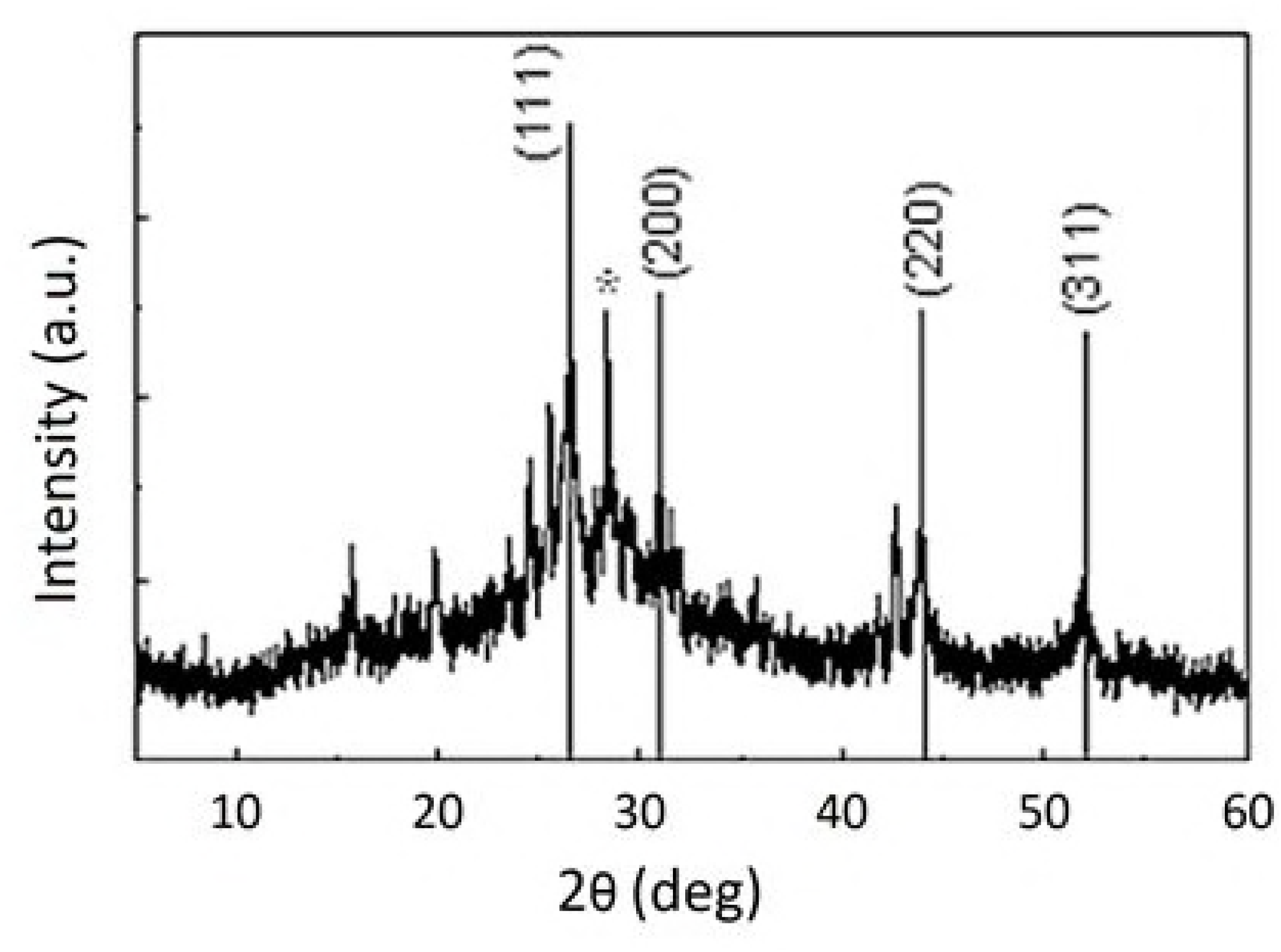
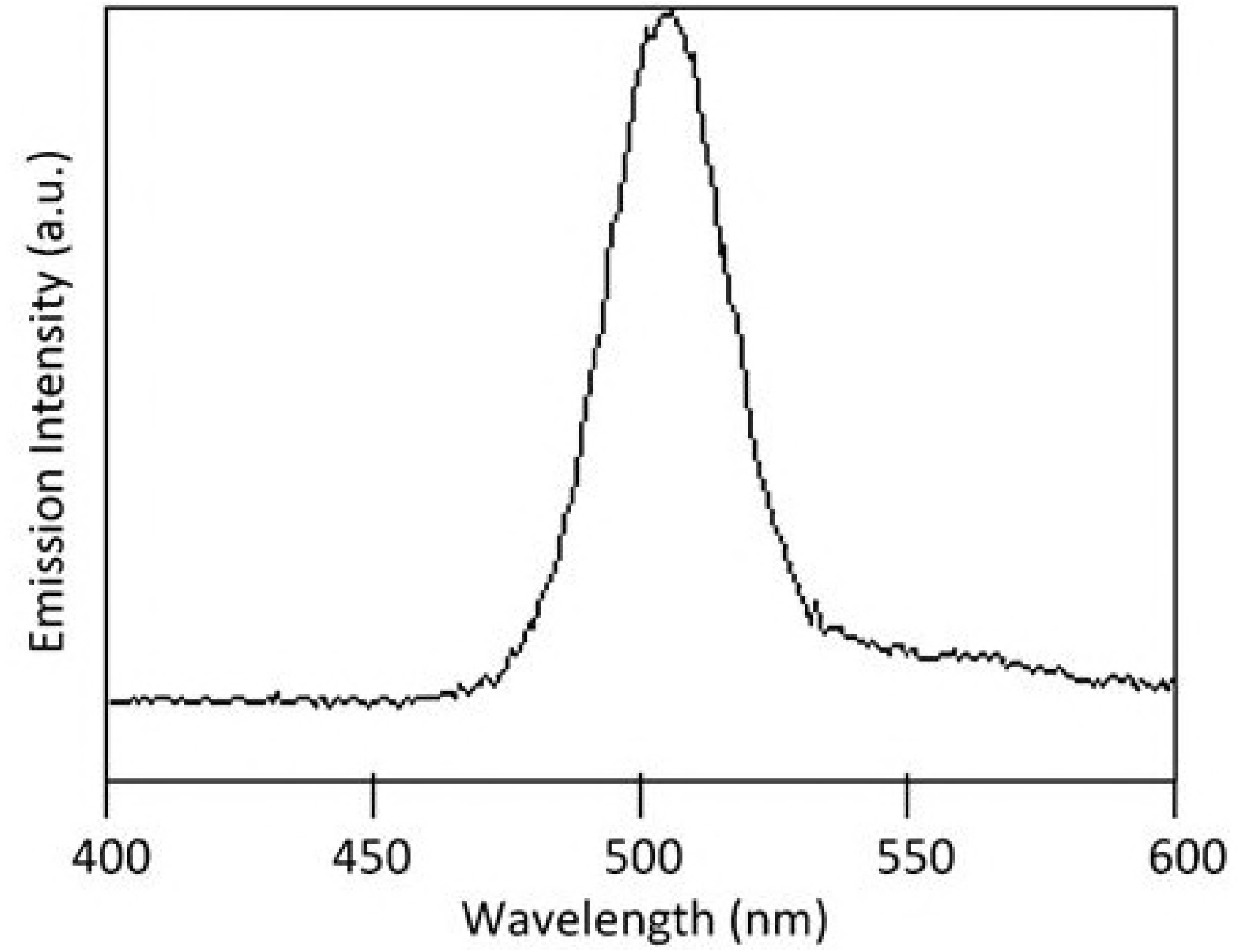
Experimental diffractogram and theoretical (vertical numbered lines) XRD for the QDs. Vertical lines correspond to cubic faces peaks. **Figure 1 -Bottom:** Emission spectrum of as prepared pegylated QDs (4nDOT500)

Cell viability results are presented in Figure 2. No significant differences between labelled (*L* set) and non-labelled (*C* set) gliomas cells were observed.

**Figure 2:**
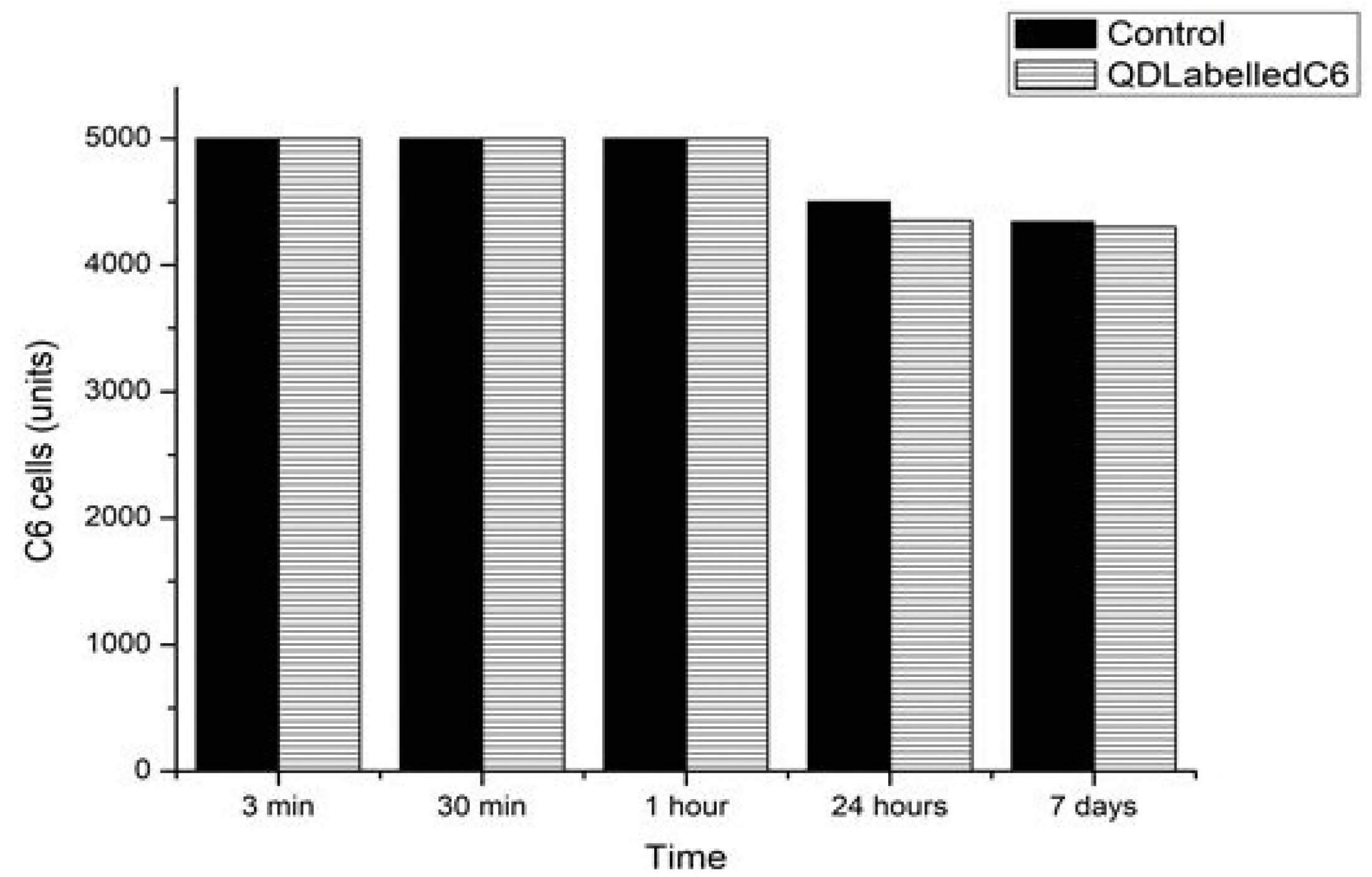
Cell viability for non-labelled (Control) C6 glioblastoma cells and for QD-labelled C6 glioma cells. The numbers are the average obtained for the 96 wells of each set.

Fluorescence image obtained by single photon Laser Scanning Confocal Microscopy for labelled C6 glioma cells, with incubation times of 3 minutes as well as 7 days are shown in Figure 3 and Figure 4, respectively.

**Figure 3:**
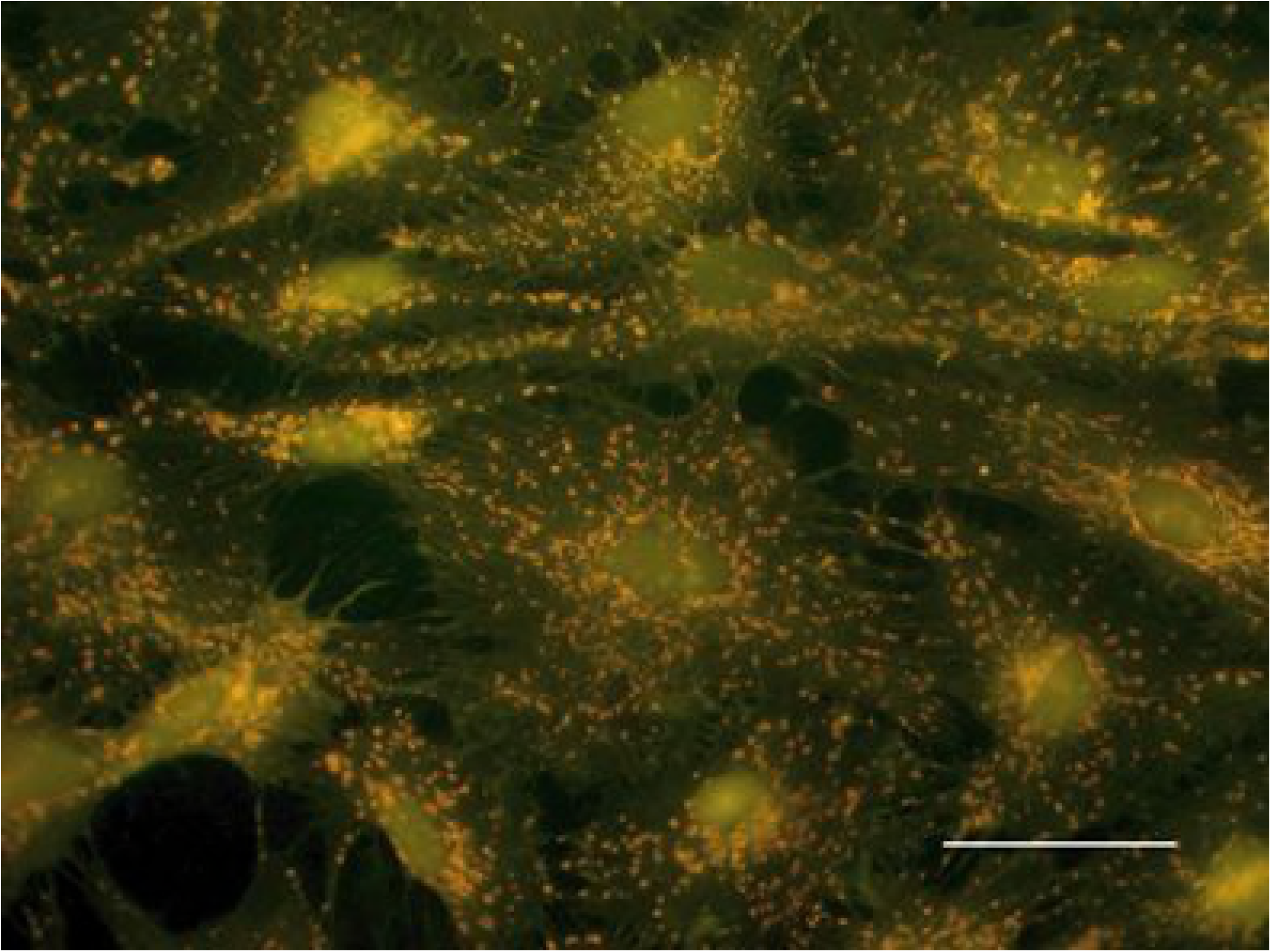
Fluorescence image of C6 glioma cells labeled with QDs (incubation time = 3 minutes), obtained by single photon laser scanning confocal microscopy. Scale bar: 25 micrometers.

**Figure 4:**
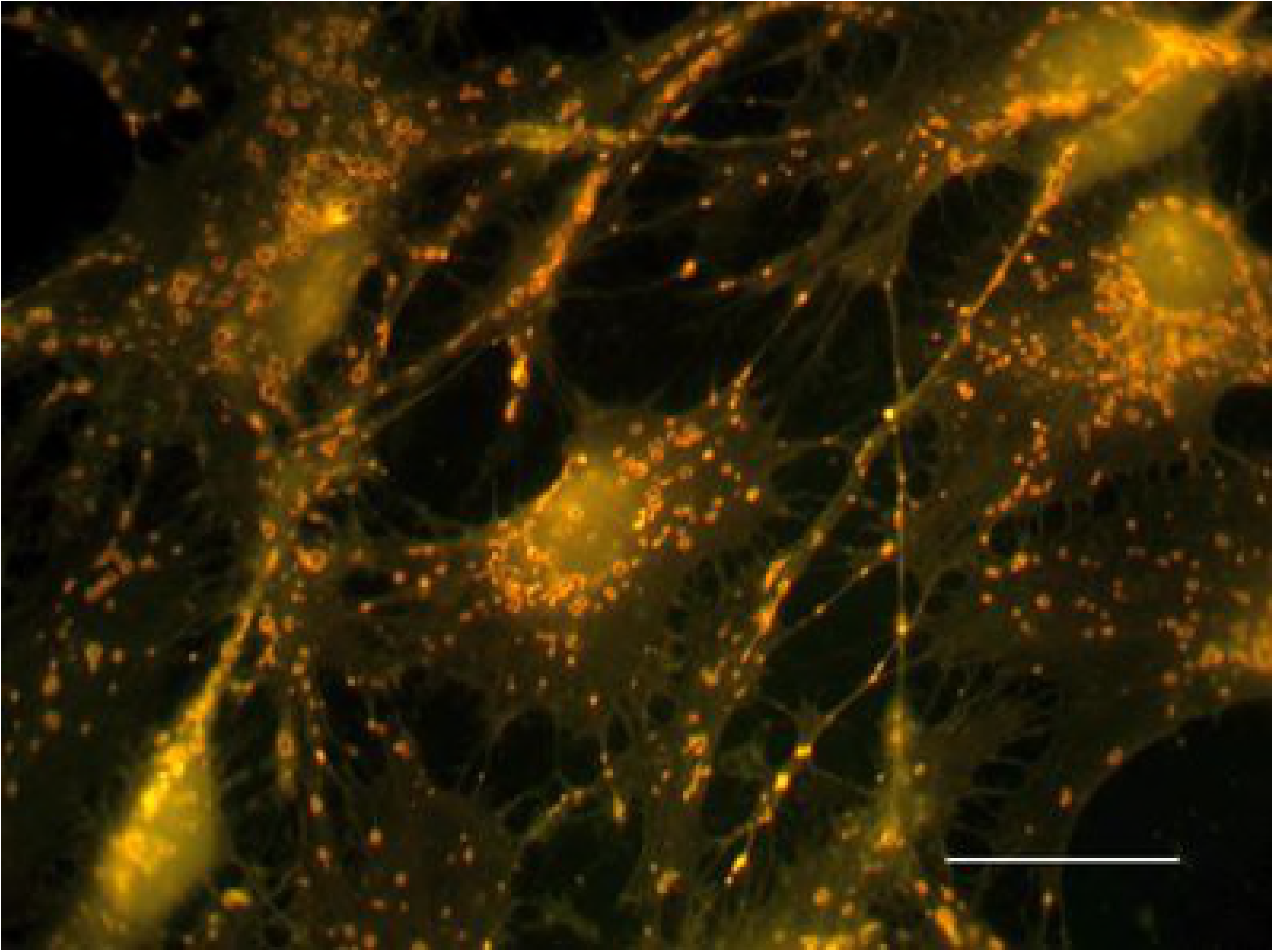
Fluorescence image of C6 glioma cells labeled with QDs (incubation time = 7 days), obtained by single photon laser scanning confocal microscopy. Scale bar: 25 micrometers.

Contrast phase image obtained for the same times, 3 minutes and 7 days, are presented in Figure 5 left and right, respectively.

**Figure 5:**
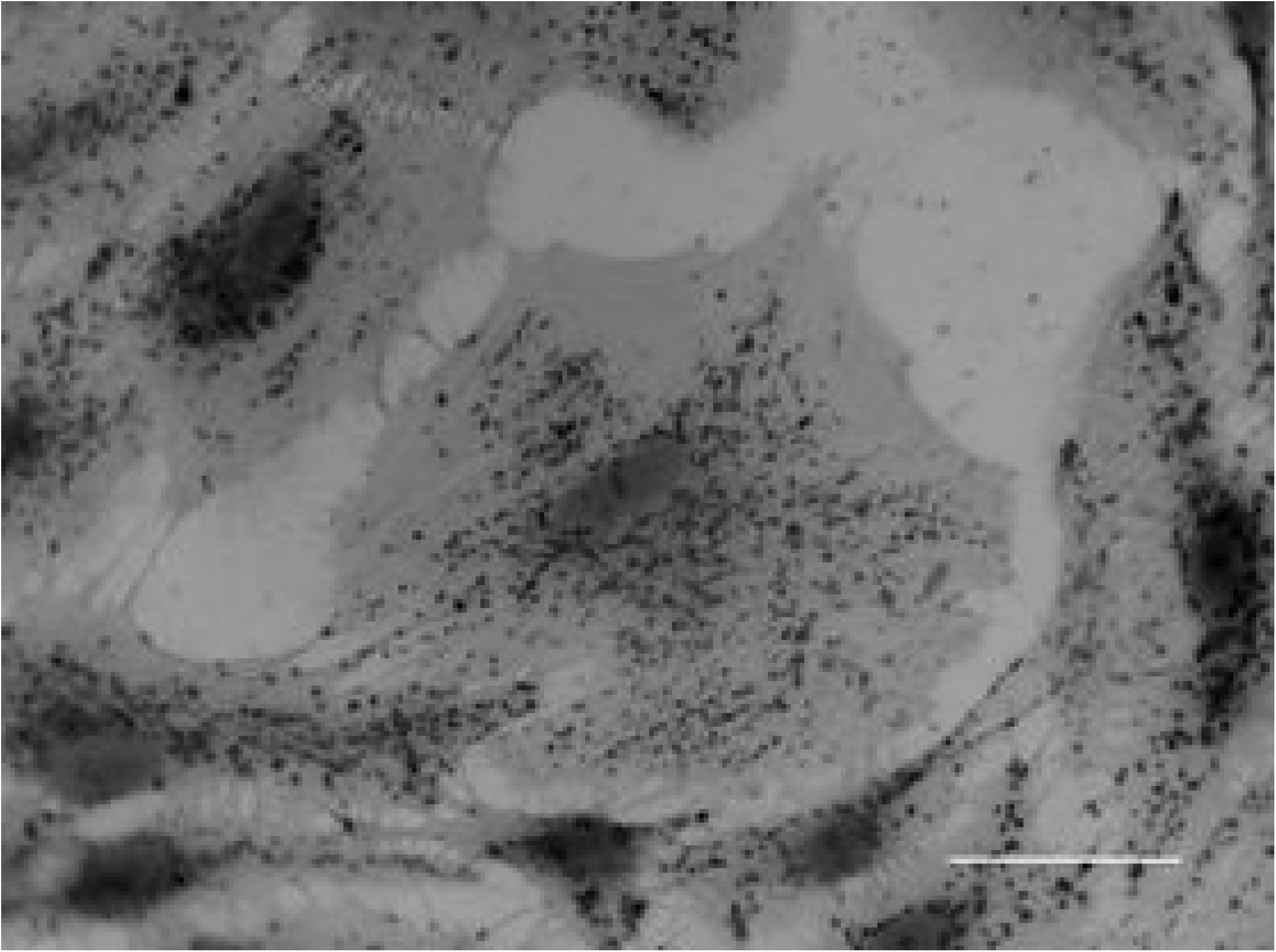
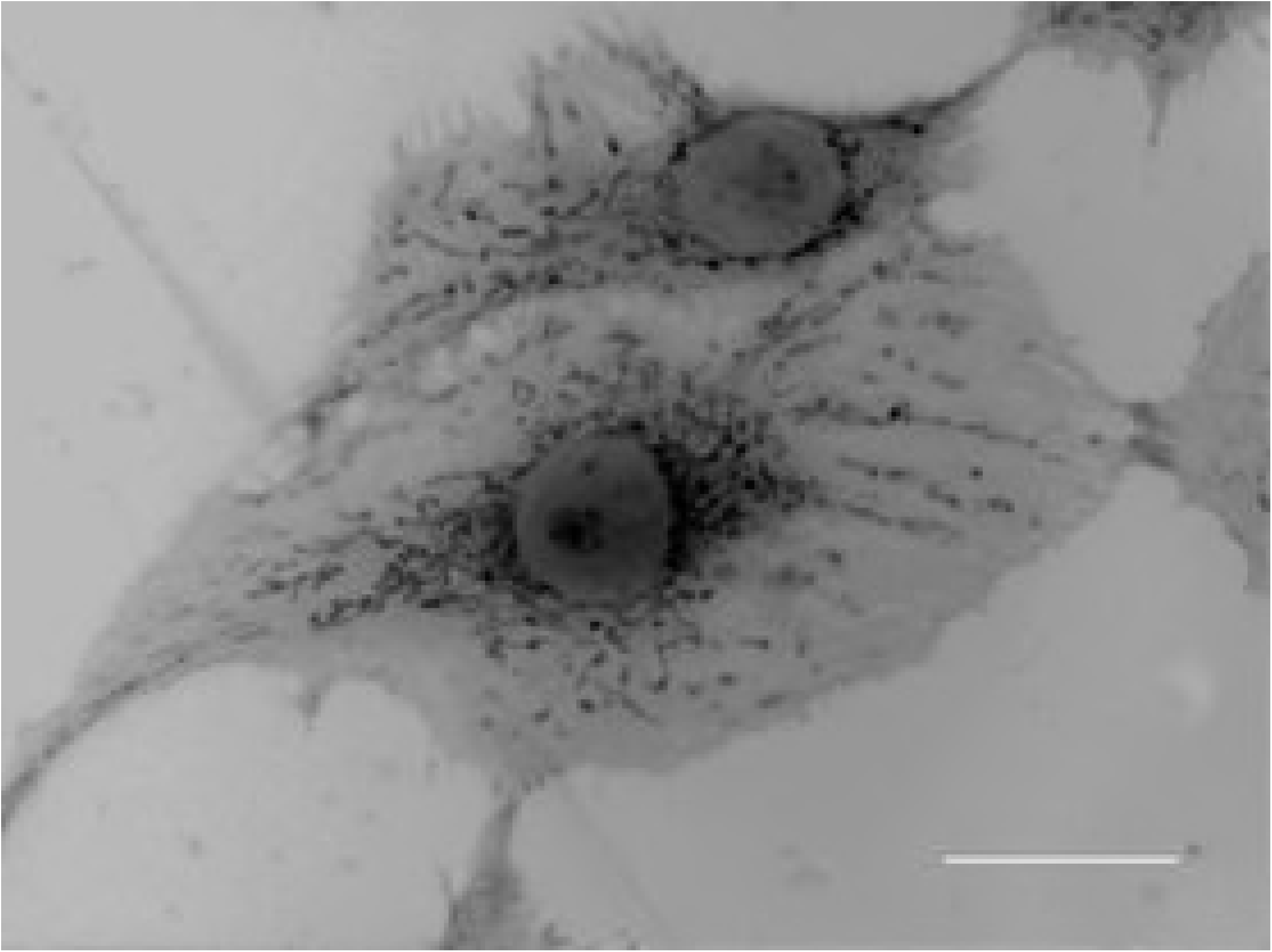
Phase contrast images of control (non-labelled) C6 glioma cells (left: 3 minutes; right: 7 days). Scale bar: 25 micrometers.

Previous *in vivo* toxicological study using same QDs, with male Wistar rats via inhalation exposure [22] have shown that the technically maximal possible concentration of free Cd^2+^ ions corresponds to 0.52 mg Cd^2+^/m³, which corresponds to a very low concentration of Cd^2+^(0.52 x10^-6^mg/ml). The results of the study indicate that there was no evidence that cadmium was translocated to the blood stream or to the central nervous system and shows that the aqueous synthesis of the QDs was very efficient, leading to a very small release rate of free ions.

## 4 CONCLUSIONS

Biocompatible, water prepared, highly fluorescent QDs were employed as probes for the long term live cell imaging of C6 glioma cells. The endocytosis of the QDs and their apparent absence of deleterious effects to the cells may be associated to the low level of free ions in the suspension containing the QDs, as well as to the efficiency of polyethylene-glycol (PEG) as functionalizing agent, which seems to bind tightly to the surface of the core/shell QDs. PEGylated QDs, *a priori*, do not have the ability to target some specific site/organelle inside the cells. In case of C6 glioma cells, the QDs were quickly internalized. In contrast, at healthy glial cells, they remained bound to cells’ surface. In both cases, presence of QDs did not negatively impact the *cell viability* and *normal cell dynamics*, which are central issues in the development of molecular imaging and drug/gene delivery systems. This study contributes as a prove of principle for a fluorescent nanoprobe-based assay, as the nanoprobes (QDs) employed have proven to be reliable for long term *in vitro* analysis and monitoring of cell mechanisms, such as migration of glioblastomas. This potentially paves the way for a better understanding of the gliomas, the most common kind of brain cancer. Further, the results allow to contribute in a positive way, to answer at least two important questions regarding long term live cell imaging [24]. The cells survived to the experiment and they could be followed in their natural behavior for a long time, with no sign of any interference nor damage.

## DECLARATIONS

The experiments were performed with line cells commercially available. All the authors consent the publication and contributed equally to this work. There are no competing interests. CNPq (Grant 405030/2015-0) and FFG (Grant 861023) partially supported this work.

